# Gelatinous fibers develop asymmetrically for posture support of bends and coils in common bean vine

**DOI:** 10.1101/2024.05.06.592750

**Authors:** Joyce G. Onyenedum, Mariane S. Sousa-Baena, Angelique A. Acevedo, Lena Hunt, Rosemary A.E. Glos, Charles T. Anderson

## Abstract

Gelatinous (G)-fibers are common in the stems of twining vines (twiners), but their role remain unclear given the lack of developmental insights. Here, we characterize the developmental anatomy of G-fiber formation in common bean stems (*Phaseolus vulgaris* L., Fabaceae). G-fibers in common bean exhibit cell wall organization comparable to other species, consisting of cellulose and Rhamnogalacturonan-I pectins, with possible traces of lignin. We show that G-fibers are absent in the actively circumnutating stems, thus these tensile fibers are not associated with the dynamic searching movements characteristics of twiners. Instead, we found that after a subtle bend or dramatic coil is formed, G-fibers form asymmetrically on the concave side of the stem for posture maintenance. Therefore, G-fibers do not drive movement, but provide support for existing bends, thus stabilizing the helical conformation of twiners around its host to avoid slippage. Finally, we present common bean as an emergent system to study twiners and growth form diversity given its easy cultivation, self-pollination, fast growth, and habit diversity arising from plant breeding, and the ability to induce habit shifts through simple modifications to light conditions.

## INTRODUCTION

All plants move their body parts to make subtle adjustments in posture, such as repositioning leaves or correcting branch angles (Hart, 1990). However, some plants move so dramatically that they become structural parasites to neighboring plants. This is the case with many vines, which survive by climbing on other plants to reach the top of the forest canopy (Darwin, 1865; Ewers et al., 2015; Stevens, 1987). Across their multiple evolutionary origins, vines have developed a wide diversity of climbing mechanisms (Acevedo-Rodríguez, 2015; Sperotto et al., 2020), including the employment of coiling stems and/or modified accessory organs such as tendrils (e.g., grape vine; *Vitis vinifera*; (Sousa-Baena et al., 2018) and gripping adventitious roots (e.g., poison ivy; *Toxicodendron radicans*).

Twining vines (hereafter “twiners”) are by far the most common type of climbing plant (Acevedo-Rodríguez, 2015; Sperotto et al., 2020) and may also be the most destructive. Some twiners wrap their main stems and branches around their host, strangling them and causing tree mortality. For example in the United States, some introduced twiners are now aggressively invasive, such as kudzu (*Pueraria montana*), bittersweet (*Celastrus orbiculatus*), and wisteria (*Wisteria sinensis* and *W. floribund*a) (Forseth and Innis, 2004; Leicht-Young et al., 2007; Trusty et al., 2007). A twiner first engages in a searching motion (“circumnutation”) to locate a support, then, by tailoring their development in response to finding a suitable host (thigmotropism) will proceed to securely coil around the support (Darwin, 1865; Sousa-Baena et al., 2021). After contact with a support, twiners must establish and maintain a tight grip on their host. So how is this helical posture maintained? Previous work has demonstrated that slight increases in the stem radius and/or development of stipules, petioles, or pulvini pinched between the vine and its support creates localized tension and generates the squeezing force that prevents stem slippage in some twiners (Isnard et al., 2009). Another possible – yet understudied – contribution to the stability of the helix is the presence of contractile Gelatinous (G)-fibers. These specialized fibers are present in several mobile plant organs, including curved tree branches (Groover, 2016), coiled tendrils of some vines (Gerbode et al., 2012; Meloche et al., 2007), and, importantly, in the stems of twiners (Bowling and Vaughn, 2009; Chery et al., 2021). The composition of G-fibers produces a strong tensile force that can bend plant organs (Gorshkova et al., 2018). These fibers contain a thickened inner wall layer (G-layer) composed of cellulose microfibrils embedded in a non-cellulosic matrix enriched in pectic Rhamnogalacturonan-I, arabinogalactan proteins, xyloglucans, and mannans (Bowling and Vaughn, 2008; Gorshkova et al., 2015; Guedes et al., 2017; Kim and Daniel, 2019; Nishikubo et al., 2007).

Previous studies have found that many twiners have G-fibers in an array of stem tissues (Bowling and Vaughn, 2009; Chery et al., 2021), however it remains unclear how these specialized fibers support the establishment of the twining habit. Here, we introduce the common bean, *Phaseolus vulgaris* L. (Fabaceae), as an emergent model system to study the relationship between G-fibers and twining growth habit. By comparing growth forms (twiner versus shrub) across and within cultivars, we determined when and where G-fibers are produced throughout the processes of stem bending, circumnutation, and coiling. We found that G-fibers are absent in circumnutating internodes and thus are not involved in generating the initial twining force. Instead, G-fibers form constitutively in stationary internodes in an asymmetric pattern, likely to help the plant maintain its posture. Finally, we argue that, given the existing diversity of growth habits produced by plant breeders and the ability to induce habit shifts with simple changes to light conditions, common bean provides a useful model for studying the development of the twining habit and growth form diversity.

## RESULTS/DISCUSSION

### Common bean displays distinct biomechanical and morphogenetic states across development

To understand the development of the twining habit in common bean, we studied internode elongation rates and timelapse movies of recombinant inbred line (RIL) L88-57 (Frahm et al., 2004), which successfully made contact with and began to climb up supports upon 9 plastochrons (22-27 days after gemination).

By carefully studying development and tracking movements through timelapse videos (**Movie 1)**, we identified five developmental stages in the establishment of the twining habit that each represent distinct biomechanical states of the stem (**Fig. 1).**

**Figure 1.**
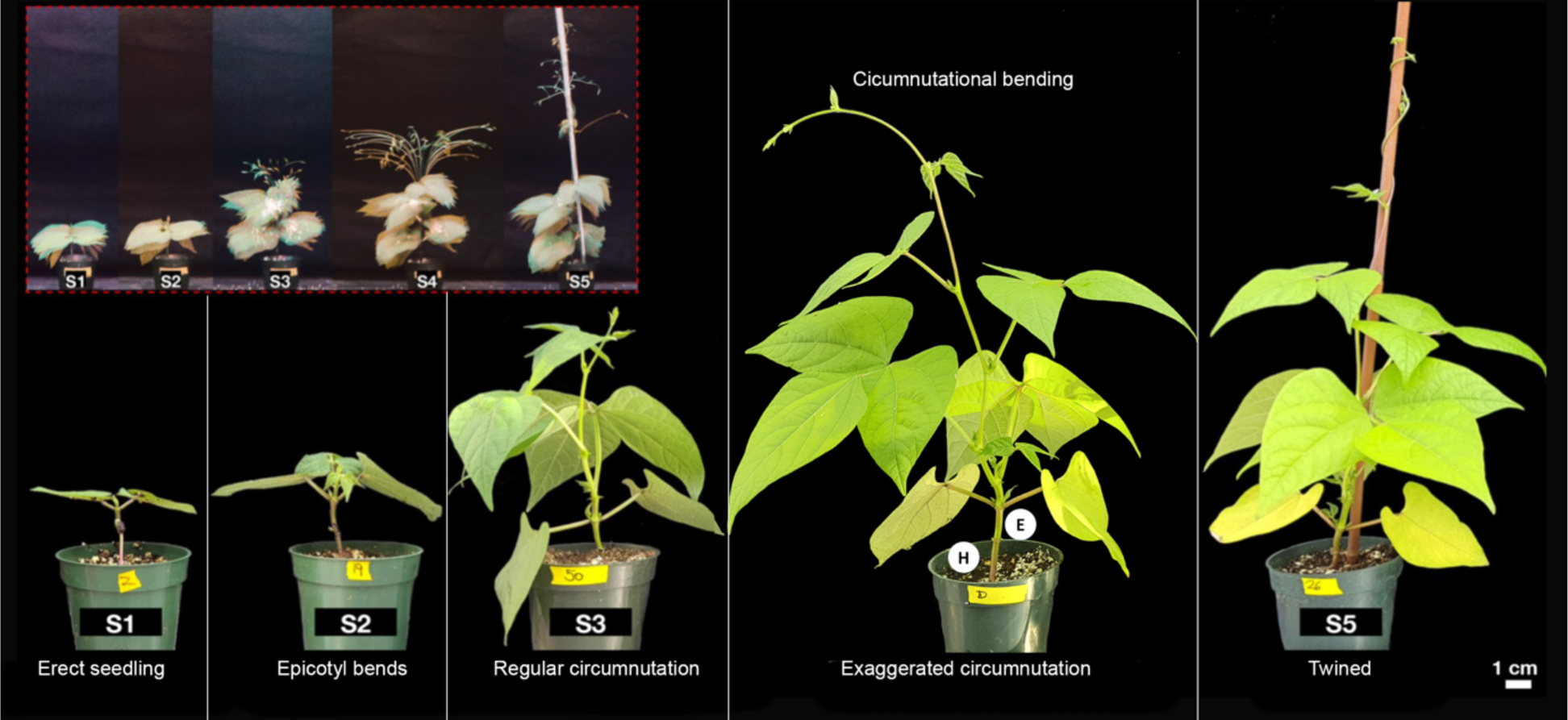
Five developmental stages (S1-S5) in the establishment of the twining habit in common bean (L88-57). Inset shows maximum intensity projections of timelapse images taken from the same individual going through the five developmental stages.

In Stage 1, plants are erect seedlings with an aboveground hypocotyl, epicotyl, and two unifoliate seedling leaves. Stage 2 plants exhibit a bent epicotyl. In Stage 3, plants undergo “regular circumnutation” (typical of all plants) with small revolutions, ≤ 9 cm in diameter. In Stage 4, the apical internodes are rapidly elongating and exhibit conspicuous circumnutational bending while performing exaggerated circumnutation with large revolutions ≥ 17 cm in diameter. Each revolution moves counterclockwise and takes 1.67 h on average, agreeing with Darwin (1875). Finally, if plants in stage 4 are given a support, within 1.33 h they complete their first gyre, thus beginning Stage 5. Guided by the leading tip, plants will continue to coil around the support until the plant dies. These five developmental stages occur at plastochron 0 (only unifoliate leaves) for Stage 1, three for Stage 2, six for Stage 3, and eight for both Stage 4 and the initiation of Stage 5.

### The common bean stem is simple, except for the presence of G-fibers

To identify which cells might be responsible for climbing, we studied the anatomy of the stem. During primary growth, the stems of common bean have the following tissues from the outside-in: a single-layer epidermis, multilayered cortex terminating in an endodermis, pericyclic fiber strands, vascular bundles in a ring, and central pith (**Fig. 2A**). Our anatomical interpretation differs from (Chernova et al., 2023) in one aspect – the innermost layer of the cortex has starch grains and can be interpreted as endodermis in bean, thus the cells just interior to this endodermis are the pericycle (Cattai and Menezes, 2010; Tamaio et al., 2009). We adopt the term “pericyclic fibers”, instead of “primary phloem fibers” as in.

**Figure 2.**
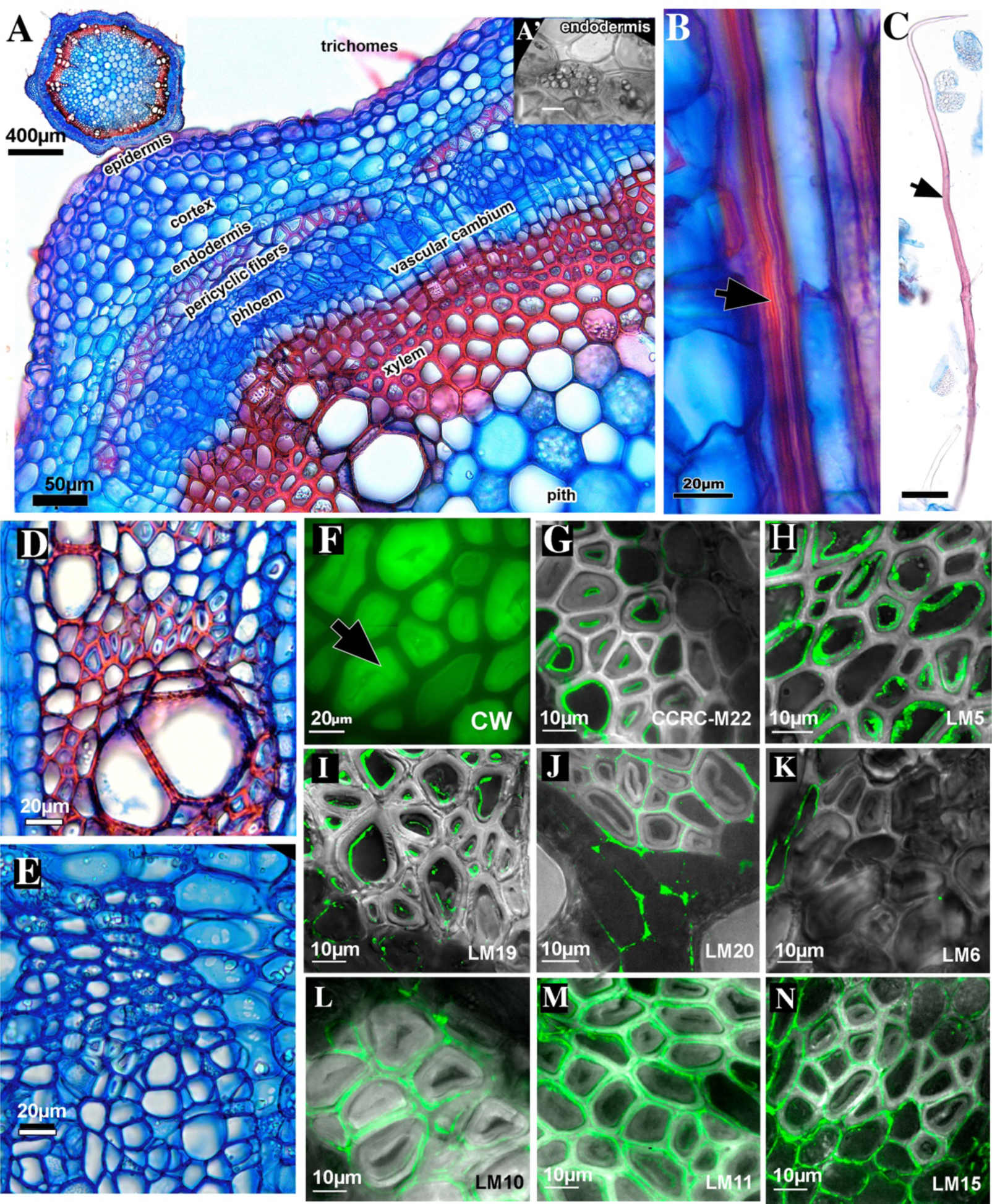
Stem anatomy and G-fiber immunolabeling in common bean. **(A)** General anatomy of common bean young internode with key tissues labeled. **A’** Inset displays starch grains delimiting the endodermis. Note the initiation of the vascular cambium, thus the tissues immediately above and below are secondary phloem and xylem, respectively. **(B)** Contracted pericyclic G-fiber **(C**) Elongated pericyclic G-fiber from a mature internode reaching ∼1.5 mm in length. **(D)** Secondary xylem and **(E)** Secondary phloem G-fibers formed later in stem development. **(F–N)** Histology and immunolabeling of cell wall epitopes in G-fibers show that G-layer is rich in cellulose and pectin. (**F)** Calcofluor White [CW] labels cellulose. (**G)** RG-I on innermost region of G-layer [CCRC-M22]. **(H)** Galactan RG-I throughout the G-layer [LM5]. **(I)** Low methyl-esterified homogalacturonan on primary cell wall/middle lamella of G-fibers [LM19]. **(J)** High methyl-esterified homogalacturonan [LM20]. (**K)** (1-5)-α-L-arabinans pectins [LM6] on inner region of G-layer **(L)** Low substituted [LM10] and **(M)** highly substituted [LM11] xylans and **(N)** XXXG motif of xyloglucan [LM15] labels the outer layers of the secondary cell walls of G-fibers.

Many vines have highly modified stem anatomies, including wholesale reorganization of the vascular tissues (Angyalossy, Pace and Lima, 2014; Cunha Neto, 2023), but common bean stems have a typical eustele and regular secondary growth. Therefore, the only clear anatomical candidate related to climbing in this species are the G-fibers; their presence has been previously reported (Chery et al 2021; Chernova et al., 2023). In young internodes, G-fibers first appear via differentiation of existing pericyclic fibers (**Fig. 2A–C**), then are later produced by the vascular cambium in the secondary xylem (**Fig. 2D**) and phloem (**Fig. 2E**) as the internode advances in maturity (see more on development in the sections below).

We confirmed that common bean has G-fibers through histological staining and immunolabeling. Staining with Calcofluor White (**Fig. 2F**) indicates that the G-layer is cellulosic. Immunolabeling assays labeled the G-layer as containing the following pectin molecules: rhamnogalacturonan-I (CCRC-M22; **Fig. 2G**), rhamnogalacturonan-I with galactan side chains (LM5; **Fig. 2H**), and low-methyl-esterified homogalacturonan pectins (LM19; **Fig. 2I**). High-methyl-esterified homogalacturonan pectins were not labeled in the G-layer, instead being labeled in the primary cell wall/middle lamella (LM20; **Fig. 2J).** (1-5)-α-L-arabinan pectin side chains were not labeled in G-fibers (LM6; **Fig. 2K**). Hemicellulose antibodies targeting xylans (LM10 and LM11; **Fig. 2L, M**) and the XXXG motif of xyloglucan (LM15; **Fig. 2N**) exclusively labeled non-G secondary wall layers. Together, these results agree with previous reports that G-fibers are characterized by the presence of a G-layer comprised mostly of cellulose and pectins.

The G-layer is often cited as being mostly cellulosic and devoid of lignin, however recent reports indicate that lignification is more widespread than previously expected (Roussel and Clair, 2015; Ghislain and Clair, 2017; Ghislain *et al*., 2019). By double staining sections with safranin and astra blue (Bukatsch, 1972) we typically found that the G-layer was blue (unlignified), whereas the outer S-layers generally stained red (lignified) (**Fig 2E).** However, we sometimes found pink G-layers (**Fig. S1A**), indicating lignification (<u>Srebotnik and Messner, 1994; Vazquez-Cooz and Meyer, 2002</u>. To further investigate these varied results, we stained sections with 2% Toluidine Blue, and found that only the outer S-layers stained blue, while the G-layer was clear (**Fig. 3B,C**), indicating the G-layer is devoid of lignin, corroborating negative results with Phloroglucinol-HCl from (Chernova *et al*., 2023). Lastly, analyzing unstained sections under UV fluorescence revealed signal in the outer S-layers and the innermost G-layer (the “Gn-layer” (Roach *et al*., 2011; Goudenhooft, Bourmaud and Baley, 2019), but not the main body of the G-layer (**Fig. S1**). This pattern of lignification matches the model in Figure 1E of <u>Ghislain and</u> <u>Clair, 2017</u>. Taken together, the G-layer in common bean is mostly devoid of lignin, with possible traces of lignin or another UV-fluorescent phenolic compound in the Gn-layer.

**Figure 3.**
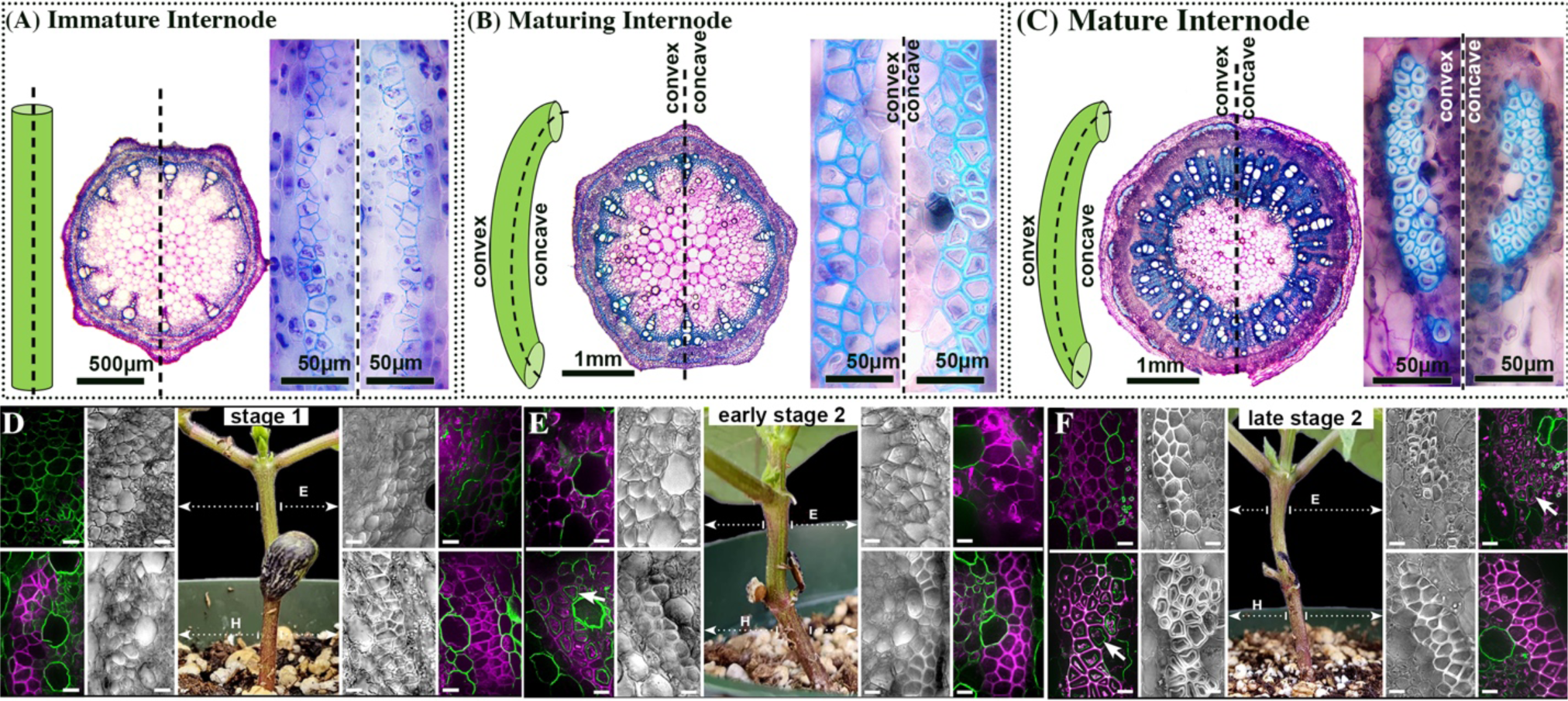
Stem shape, cross section, and differentiation of G-fibers in young and maturing internodes. **(A)** Immature internodes emerge straight; pericyclic fibers do not have G-layers (left and right insets). **(B)** Maturing internodes are bent; pericyclic fibers differentiate into G-fibers only on the concave side (right insert). **(C)** Mature internodes are bent; pericyclic G-fibers are now present on both concave and convex sides (insets) (**D–F)** Developmental Stages 1-2, showing how pericyclic G-fibers form after a slight bend. **(D)** Stage 1, the epicotyl is straight and devoid of G-layers. **(E)** Early Stage 2, a subtle bend in the epicotyl appears. No G-fibers are present in the epicotyl. (**F)** Late Stage 2, the subtle bend is reinforced by asymmetric production of G-fibers on the concave side of the epicotyl. D**–**F, Scale bars=10 µm. E, Epicotyl; H, Hypocotyl.

### Asymmetric G-fiber development reinforces subtle bends in each internode

To understand the relationship between G-fibers and stem curvature, we began by identifying when and where G-fibers form during the maturation of a single internode. Immature internodes emerge straight and all pericyclic fibers sheathing the stele have thin, lignified secondary walls without G-layers **(Fig. 3A&D**). Shortly thereafter, a subtle bend in the internode is observable, then G-fibers form asymmetrically **(Fig. 3B, E-F)**. The pericyclic fibers on the concave side differentiate by developing G-layers, whereas the pericyclic fibers on the convex side do not (**Fig. 3B, E-F)**. This pattern is analogous to the tension-opposite wood division in trees (Groover, 2016), and the asymmetric contraction of G-fibers along the dorsiventral axis of cucumber tendrils (Gerbode 2012). Our finding, that G-fibers appear only after a bend is present, demonstrates that G-fibers reinforce existing bends rather than creating them.

As an internode matures, the existing G-fibers on the concave side progressively mature with thicker G-layers, while the ordinary fibers on the convex side generate G-layers (**Fig. 3C**). Concomitantly, the vascular cambium produces G-fibers mainly in the secondary xylem (**Fig. 2D)**, and rarely in the secondary phloem (**Fig. 2E)** in a seemingly haphazard fashion. We found secondary G-fibers only in the hypocotyl and epicotyl of mature, twined plants, suggesting that pericyclic G-fibers, which are present in young internodes, are likely more important in the initial establishment of the twining habit.

### G-fiber distribution along the plant axis is linked to posture maintenance

Having found that pericyclic G-fibers are associated with the concave side of a localized, recent bend, we next sought to understand how the asymmetric production of pericyclic G-fibers relates to overall plant habit in twiners vs. shrubs. Towards this aim, we first tracked the positions of G-fibers throughout the five developmental stages of the twining habit, as described above. G-fibers are absent in newly emergent seedlings (Stage 1, **Fig. 3D**). The first sign of pericyclic G-fibers is in Stage 2, where they are present exclusively on one side of the hypocotyl, while the epicotyl is slightly bent. This bend is formed before G-fibers are present (**Fig. 3E)** but is quickly followed by the production of G-fibers on the concave side of the curve; G-fibers are absent on the convex side (**Fig. 3F)**. In Stage 2, G-fibers are present along the shaft in an alternating pattern from hypocotyl to epicotyl (**Fig. 3F)**. During both Stages 3 (regular circumnutation) and 4 (exaggerated circumnutation), twiners are under a biomechanical state where the base of the plant remains stable and erect while the apical internodes are dynamically moving. Along the basal erect internodes (hypocotyl to internode 2), the alternating pattern of G-fibers is maintained, while the dynamic apical internodes are devoid of G-fibers (**Fig. S2**). Upon attachment to a support stake (Stage 5), the alternating pattern of G-fibers becomes disrupted **(Fig. 4A)**. In stage 5, twined internodes (3 to 5) have G-fibers located exclusively on the inside of the coils, following the path of a right-handed helix. The apical internodes (6-9) do not possess G-fibers yet as they are immature and only lightly attached. Taken together, G-fibers are not present in actively mobile internodes, thus are not associated with generating the circumnutational force. Instead, G-fibers form after an internode becomes stationary, securing localized bends and stabilizing coiled internodes to avoid stem slippage.

**Figure 4.**
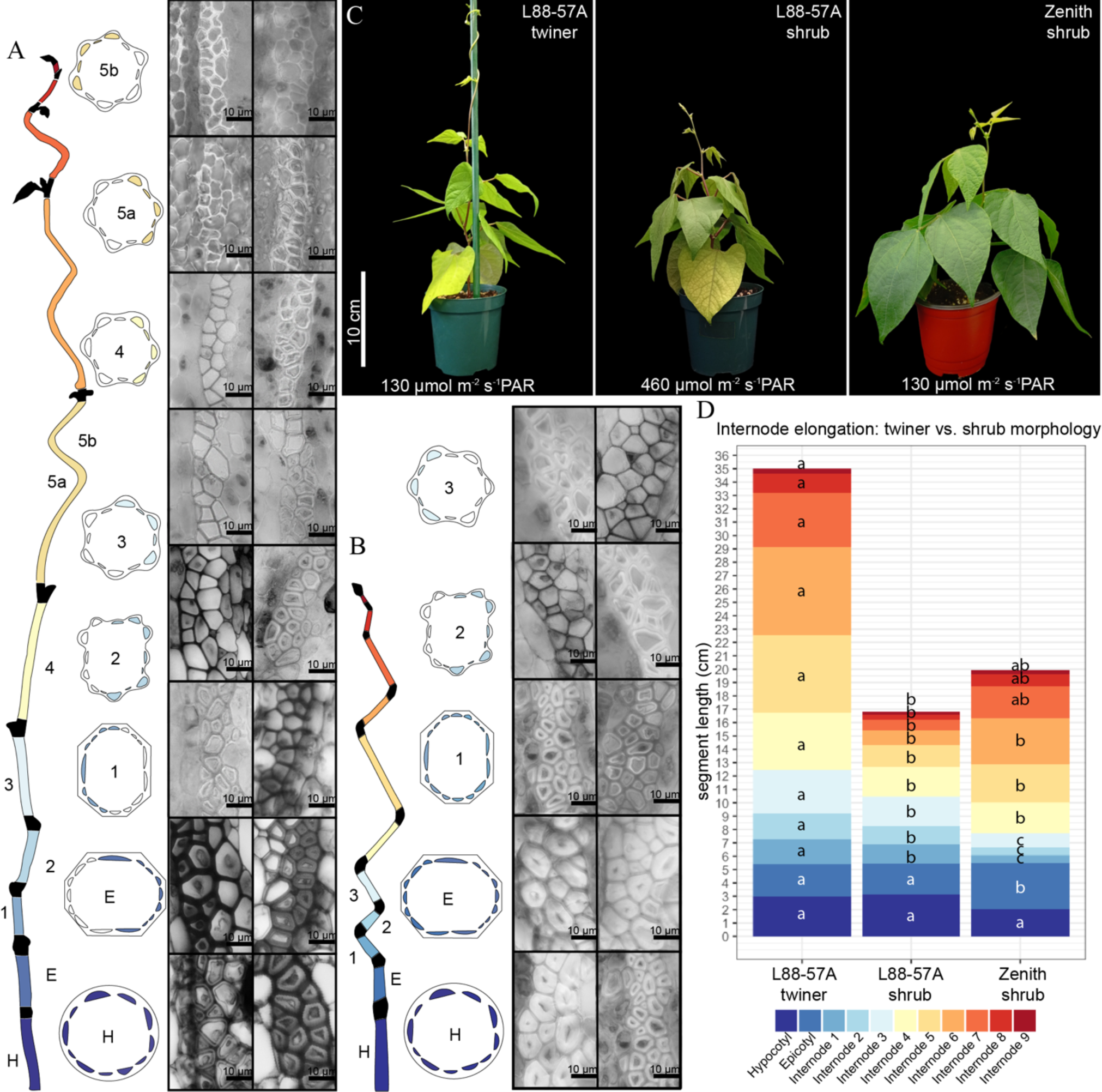
G-fiber distribution in twiner vs. shrub common bean plants. **(A)** Unraveled L88-57 twiners reveal a mature hypocotyl with all pericyclic G-fibers differentiated as G-fibers, followed by asymmetrical G-fiber differentiation occurring on alternating sides of the stem from epicotyl to internode 2. In internodes 3 to 5, G-fibers are localized along the inside of the stem. **(B**) L88-57 shrub, pericyclic G-fibers sheath the entire stem circumference in hypocotyl, epicotyl, and internode 1. Internodes 2 and 3 are relatively immature given asymmetric G-fiber distribution. **(C)** Habit diversity in this study; L88-57, grown as a twiner in low light conditions; L88-57, grown as a shrub under high light; and, ‘Zenith’, grown as a shrub under low light. **(D)** Internode elongation comparison between L88-57 twiner, L88-57 shrub, and ‘Zenith’ shrub at plastochron 9. Different letters indicate significant difference (p<0.05). by one-way ANOVA and TukeyHSD post hoc test.

To further analyze the function of G-fibers in posture maintenance, we compared G-fiber distribution in twiners and shrubs (**Fig. 4; Fig. S3-6**). Many common bean cultivars adjust their growth habit according to light conditions (Van Dobben, Van Ast and Corré, 1981). Shrubs were grown in two ways: (1) Shrub cultivar ‘Zenith’ under low light (130 µmol/m^2^/s), and (2) Typically twining L88-57 plants under high light conditions. All twiners in this study were grown under low light (130 µmol/m^2^/s). All ‘Zenith’ and L88-57 shrubs grew erect and were composed of truncated internodes in comparison to L88-57 twiners (**Fig. 4D**). The total height of ‘Zenith’ (mean = 21.4 cm, 3.2 cm, n=8) and L88-57 shrubs (mean=18.7 cm, SD= 2.1 cm, n=18) was not significantly different (Welch Two sample T-test, t = -2.1778, df = 9.7015, p-value = 0.05526), but individual internodes elongated differentially (**Fig. 4D**). Developmental progression of twiners and shrubs was similar until plants reached plastochron seven, at which point internode 3 elongates more in twiners than shrubs (**Fig. S7**). Like twiners, each internode in shrubs had a subtle curvature corresponding to a concave and convex side (**Fig. S8**). Both shrub phenotypes possessed G-fibers encircling the stem circumference in not just the hypocotyl and epicotyl but also internode 1, therefore shrubs had more mature basal internodes than twiners **(Fig. 4A, B)**. G-fibers were asymmetrically produced on alternating sides of the stem from internodes 2-4 in shrubs, (**Fig. 4B, Fig. S3-6)**, like the erect basal internodes of twiners.

Given our findings that (1) G-fibers are present in twiners and shrubs and (2) G-fibers form prior to any sign of habit differentiation, we conclude that G-fibers are not exclusively associated with the twining habit, but instead form constitutively within the species. Therefore, G-fibers are useful for stabilizing multiple types of localized bends and coils in common bean.

## CONCLUSIONS

In this study, we characterized the developmental anatomy of G-fiber formation in the stems of common bean. Within an individual internode, G-fibers first formed asymmetrically via differentiation of pericyclic fibers on the concave side of an existing bend. Subsequently, G-fibers arose from the vascular cambium mostly in the xylem in a haphazard fashion, and seldom in the phloem. G-fibers in common bean exhibited cell wall organization comparable to those reported in other species, consisting of cellulose and Rhamnogalacturonan-I pectins, with possible traces of lignin or another phenolic compound. By tracking the position of pericyclic G-fibers throughout the five stages in the establishment of the twining habit, we found that G-fibers are absent in immature and/or actively circumnutating internodes, indicating that they are not involved in rapid dynamic movements. Instead, G-fibers formed constitutively in stationary internodes, where they develop in an alternating asymmetric pattern likely for posture maintenance of erect internodes in twiner and shrub phenotypes, and in the inner side of twined internodes to stabilize the stem helical conformation. Given the combination of (1) fast growth rate, (2) the habit diversity already made through plant breeding, and (3) the inducibility of habit shifts with simple light manipulations, we propose common bean as an emergent model species to study the development of the twining habit and growth form diversity.

## METHODS

### Plant cultivation

*Phaseolus vulgaris* L., RIL L88-57 (Frahm *et al*., 2004) and upright shrub cultivar ‘Zenith’ black bean (Reg. No. CV307, PI 673047; (Kelly *et al*., 2015) were generously provided by Johnathan Lynch (The Pennsylvania State University) and Dr. James Kelly from Michigan State University, respectively. The specific L88-57 line was “A” in Dr. Lynch’s laboratory. L88-57 is the progeny of a cross between a type II upright short vine with limited branches and a type III vine with opportunistic branching (Dr. James D. Kelly, personal communication; Frahm et al., 2004). Plants were germinated at Cornell University and New York University under the following conditions: seeds were primed in a solution of 20% PEG 4000 in the dark for 24 h, subsequently rinsed thoroughly and incubated for 48 h in Petri dishes on water-saturated filter paper at room temperature on the laboratory bench.

L88-57 A were grown under two conditions to either induce climbing (low-light) or induce a shrub phenotype (high-light condition). The low light conditions include 12 hrs light with 130 µmol/m^2^/s PAR, a 30-minute ramp down, and 12 h dark, followed by a 30-minute ramp up, with 50% humidity in Environmental Growth Chamber (EGC) Model GR48 and SLR-90. Additional low light plants were grown under the following conditions: 14 h light with 130 µmol/m^2^/s PAR, 10 h dark, with 50% humidity in the Cornell University Agricultural Experiment Station Guterman greenhouses (Ithaca, NY). High light conditions include 12 hrs light at 460 µmol/m^2^/s PAR with a 30-minute ramp down, 12 hrs dark, followed by a 30-minute ramp up, with 50% humidity in EGC Growth Chamber models. ‘Zenith’ plants were grown under similar low light conditions as described above, except with a 16 h light period and 8 h dark period in the EGC Growth Chamber models GR48 and SLR-90. L88-57A plants were grown using LM-111 All Purpose Soil Mix, while ‘Zenith’ was grown using Glee All Purpose Potting Mix. Fertilizer, 75ppm of Jacks 15-5-15, was applied as needed. Internode elongation was obtained for plants at plastochrons 0 to 9. A new plastochron is designated when the midleaf of the apical trifoliate reaches 0.3 cm length.

### Timelapse videography

Raspberry Pi III (https://www.raspberrypi.com/) computers were programmed with Raspberry Pi Camera V2 to image 1 frame every 20 minutes for 3 weeks. Plants were placed in front of a black velvet backdrop. We measured the diameter of the circumnutation revolution (roughly circular trajectory described by the shoot tip) from maximum intensity projections of timelapse images. Three timelapse videos were used to obtain the diameter of circumnutation, the timing to complete one gyre, and the timing of one circumnutational revolution.

### Developmental Anatomy

The five developmental stages (1. Erect seedling, 2. Epicotyl bends, 3. Regular circumnutation, 4. Exaggerated circumnutation, 5. Twined). At each developmental stage, each internode was sectioned and analyzed according to the following protocol: stem sections were fixed in formalin acetic acid-alcohol (3 days), and then permanently stored in 70% ethanol. Following transfer to ethanol, stem sections were taken by hand with an ASTRA razor blade (model: ASTRA01) from the middle of each internode. For the hypocotyls, sections were taken closer toward the epicotyl to obtain sections with stem-like, rather than root-like, anatomy. To differentiate lignified from non-lignified cellulosic cell walls, sections were double stained with safranin and astra blue (Bukatsch, 1972). Specifically, we used 1% safranin-O (Fisher, 50 180 6544) in 50% ethanol and 1% astra blue (Santa Cruz Biotechnology, sc-214558) in 50% ethanol in a 9:1 ratio. Sections were also separately stained with 2% Toluidine Blue in water as in <u>Pradhan Mitra and</u> <u>Loqué, 2014)</u>. Stained sections were mounted in glycerol, and then imaged under light microscopy using an Olympus BH2 microscope and AmScope MU 1000 digital camera (https://amscope.com/). Additionally, unstained sections were imaged under UV light (405 nm excitation, 410-450 emission) using a Leica Stellaris 5 Confocal. For developmental stages 1–3, we also detected developing G-fibers with immunolocalization using an anti-RG-I primary antibody (LM5) and Alexa 488 Goat anti-Rat IgG Fab fragment secondary antibody (see Methods: Immunolocalization).

To visualize the localization of cellulose with fluorescence microscopy, we stained sections with Calcofluor White (Krackeler Scientific, 45-18909-100ML-F-EA) and imaged according to the following procedure: Hand sections were stained at RT in darkness for 5 min in aqueous 0.01% Calcofluor White with a few drops of ∼1 N sodium hydroxide (Fisher, BP359 500) added. Stained sections were rinsed in DI water and mounted in glycerol on glass slides. Slides were imaged with a 405 nm excitation laser and 405/60 nm emission filter on an Andor Revolution Spinning Disk Confocal Microscope with Inverted Olympus IX-83 microscope at the Cornell Institute of Biotechnology’s Imaging Facility.

### Cell wall Immunohistochemistry

We employed immunolocalization to label the following cell wall epitopes: homogalacturonan pectins (LM19,LM20; (Verhertbruggen *et al*., 2009); rhamnogalacturonan-I (LM5(Jones, Seymour and Knox, 1997, p. 199) arabinan side chain (LM6;(Willats, Marcus and Knox, 1998), xylans (LM10, LM11; (McCartney, Marcus and Knox, 2005), xyloglucan (LM15; (Marcus *et al*., 2008). Cross sections were hand sliced with Astra Razor blades, blocked in 1 x PBS + 5% (w/v) non-fat powdered milk (Carnation) in 1.7 mL Eppendorf tubes for 40 minutes, then washed three times with 1x PBS. Sections were incubated in primary antibody diluted 1:10 in 1x PBS + 5% Carnation milk powder for 90 min, then washed three times in 1x PBS. Following this, sections were incubated in secondary antibody diluted 1:100 in 1 x PBS + 5% non-fat powdered milk for 90 min, then washed three times in 1x PBS; Rhodamine Red™-X (RRX) Goat Anti-Mouse IgG Fab fragment for CCRC-M22 primary antibody, and Alexa 488 Goat anti-Rat IgG Fab fragment for all other primary antibodies. Negative controls only contained the secondary antibodies to observe non-specific fluorescence. Confocal images were collected on a Zeiss Axio Observer microscope with a Yokogawa CSU-X1 spinning disk head and a 63X or 100X 1.4 NA oil immersion objective. A 488 nm excitation laser and a 525/550 nm emission filter were used for the detection of epitopes tagged with Alexa 488 Goat anti-Rat IgG Fab fragment and a 561 nm excitation laser and a 617/673 nm emission filter was used to image epitopes tagged with Rhodamine Red™ X (RRX) Goat Anti-Mouse IgG Fab fragment. For each antibody, two experiments were performed; each experiment comprised three cross sections of hypocotyls from three control plants.

## ACKNOWLEDGEMENTS AND FUNDING

The authors thank the Cornell Institute of Biotechnology’s Imaging Facility for access to the Andor Revolution Spinning Disk Confocal Microscope, funded by NIH 1S10OD010605. This work was mainly supported by NSF CAREER award #2401675 to J.G.O. Parts of the Growth & Development and Immunolocalization experiments were supported as part of The Center for Lignocellulose Structure and Formation, an Energy Frontier Research Center funded by the U.S. Department of Energy (DOE), Office of Science, Basic Energy Sciences (BES), under Award # DE-SC0001090 to C.T.A.

## AUTHOR CONTRIBUTIONS

J.G.O. and C.T.A conceived the project. J.G.O, M.S.S.B., A.A.A., R.A.E.G., and L.H. designed and performed the experiment, and analyzed data. J.G.O and A.A.A. wrote scripts. J.G.O and M.S.S.B wrote the initial article. All authors contributed to the final manuscript.

---

**Figure S1:**
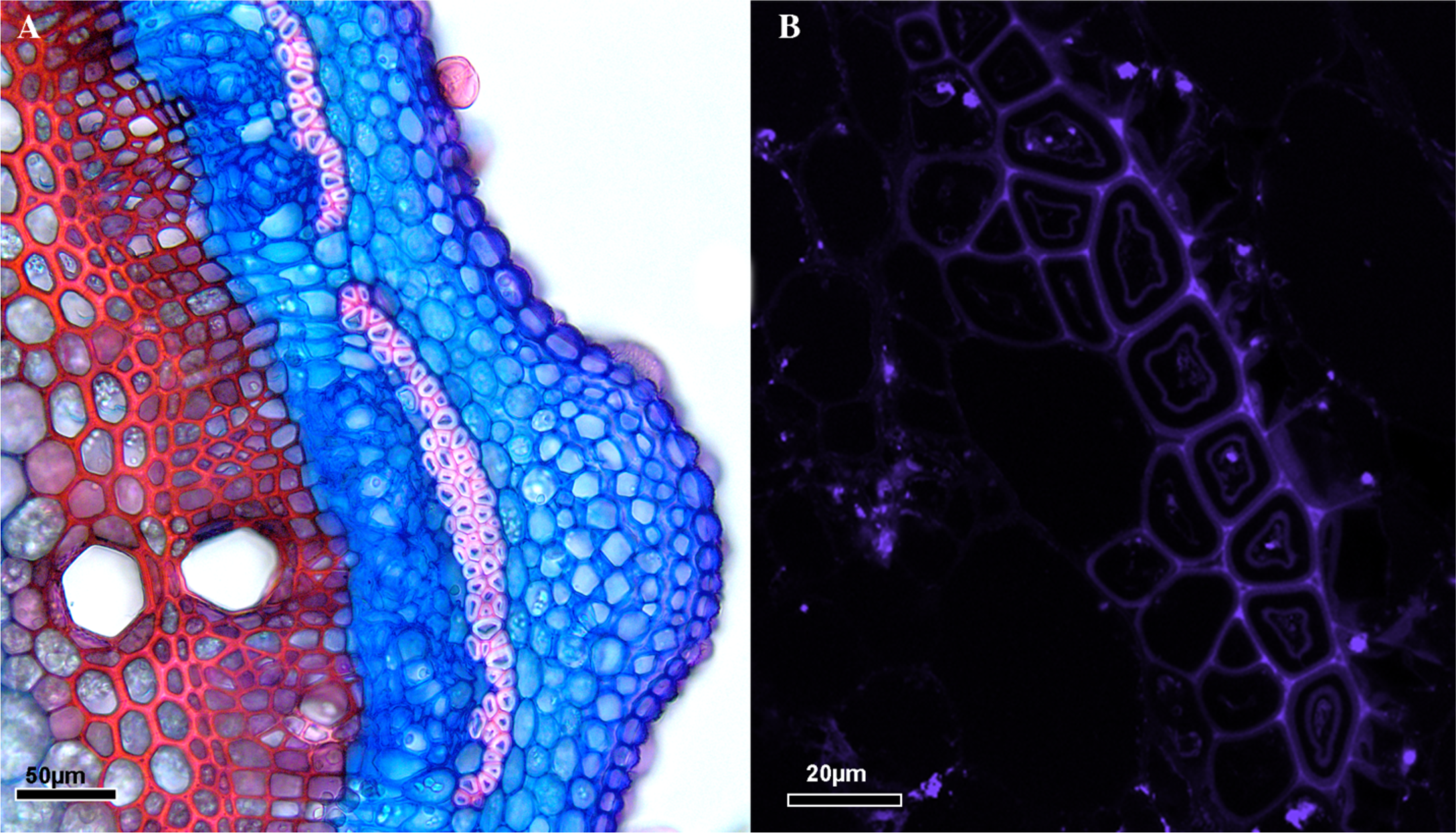
Differential results concerning lignification of the G-layer in L88-57 Twiner. **(A)**> Safranin + Astra Blue double staining sometimes yields pink G-layers indicating lignification. **(B)** UV autofluorescence indicates lignification only on the S-layers and innermost G-layer fluoresces; not the body of the G-layer itself.

**Figure S2:**
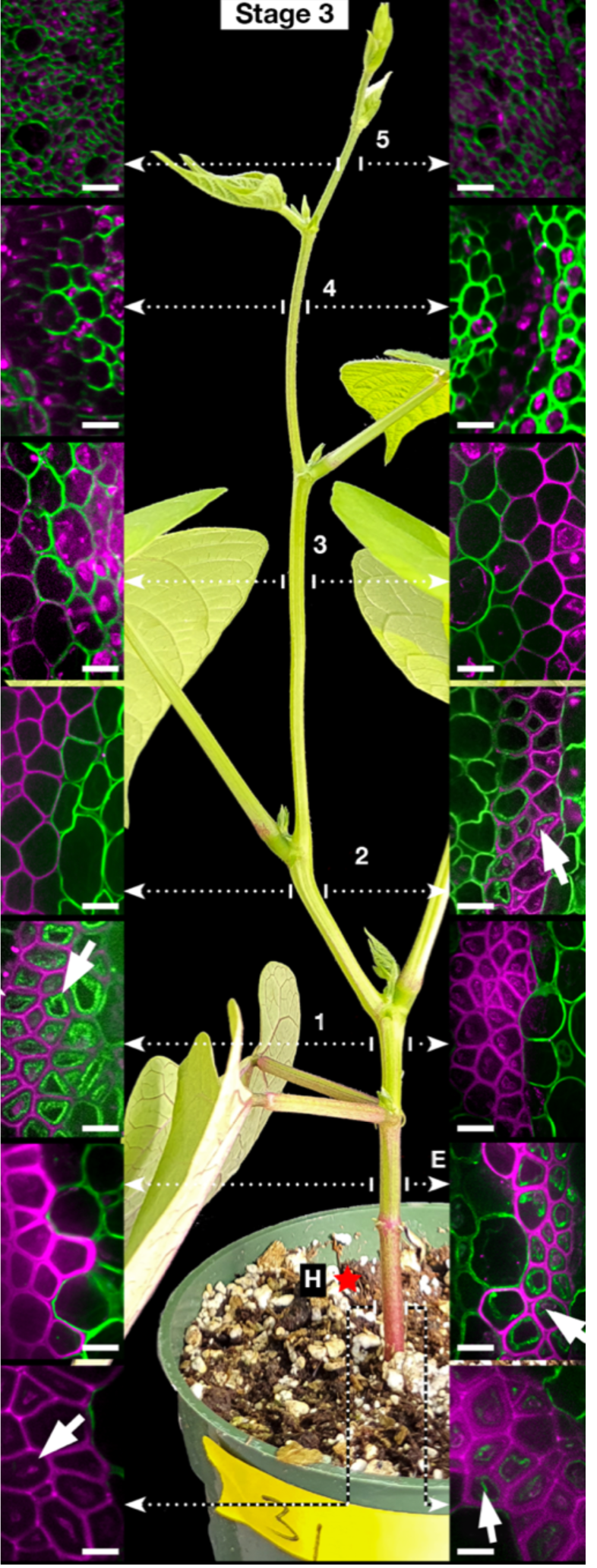
Pericyclic G-fiber distribution in stages 3 (regular circumnutation) in L88-57 Twiner. Pericyclic fibers differentiate asymmetrical and on alternating sides of the stem from hypocotyl to internode 2. Note that hypocotyl has G-fibers on both sides of the stem: the left image are the original G-fibers which have matured, and the right-hand image are newly developed G-fibers with thin G-layers (see section “Asymmetric G-fiber development reinforce subtle bends in each internode” for further discussion). G-fibers are absent from internode 3 upwards because these internodes are immature and undergoing regular circumnutation. White arrows indicate the G-layers. Scale bars=10 µm. E, Epicotyl; H, Hypocotyl.

**Figure S3:**
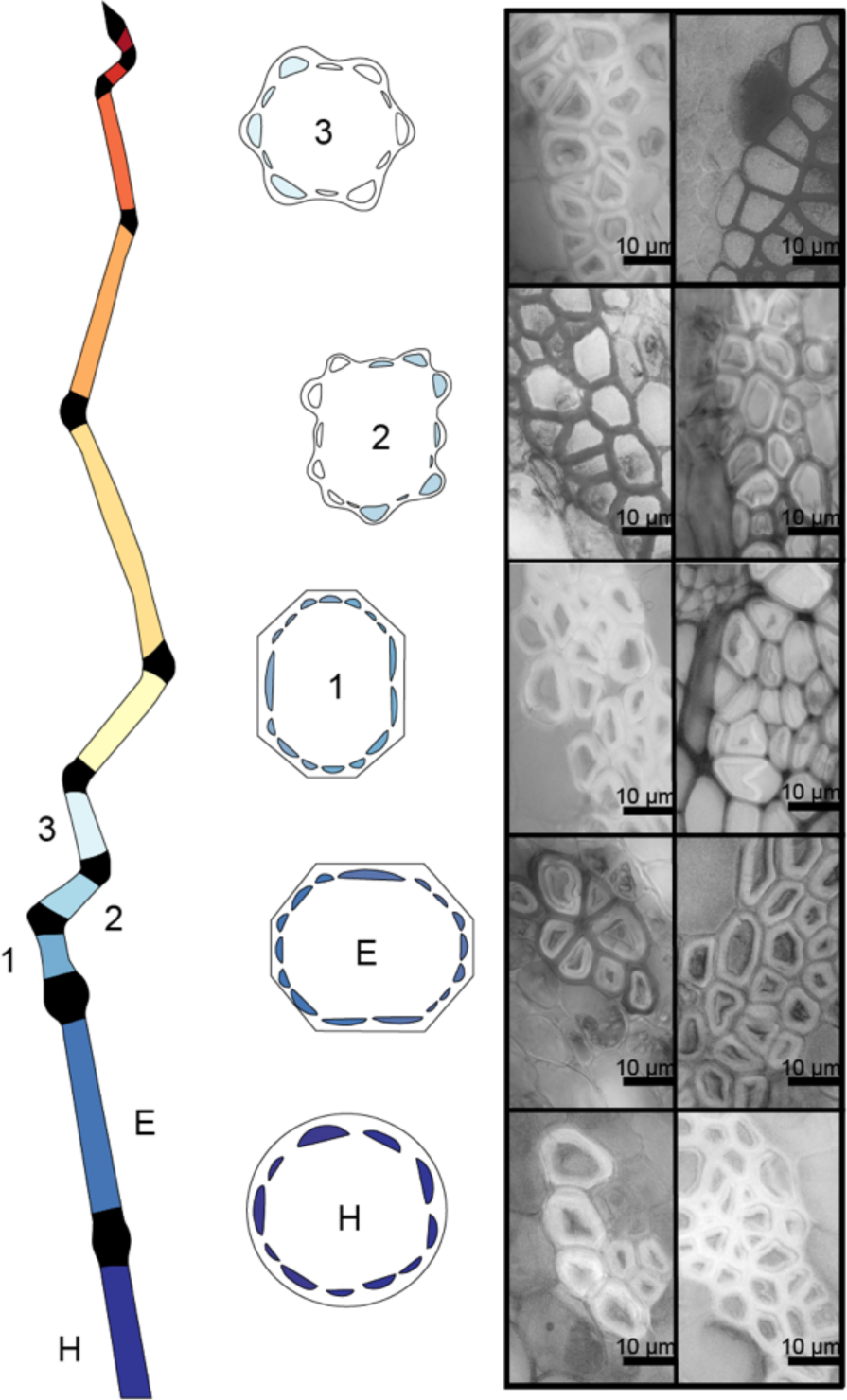
‘Zenith’ pericyclic G-fibers sheath the entire stem circumference in hypocotyl, epicotyl, and internode (like L88-57 shrubs), thus shrub phenotype have more mature basal internodes than twiners. Internodes 2 and 3 are relatively immature evident by the asymmetric G-fiber distribution.

**Figure S4:**
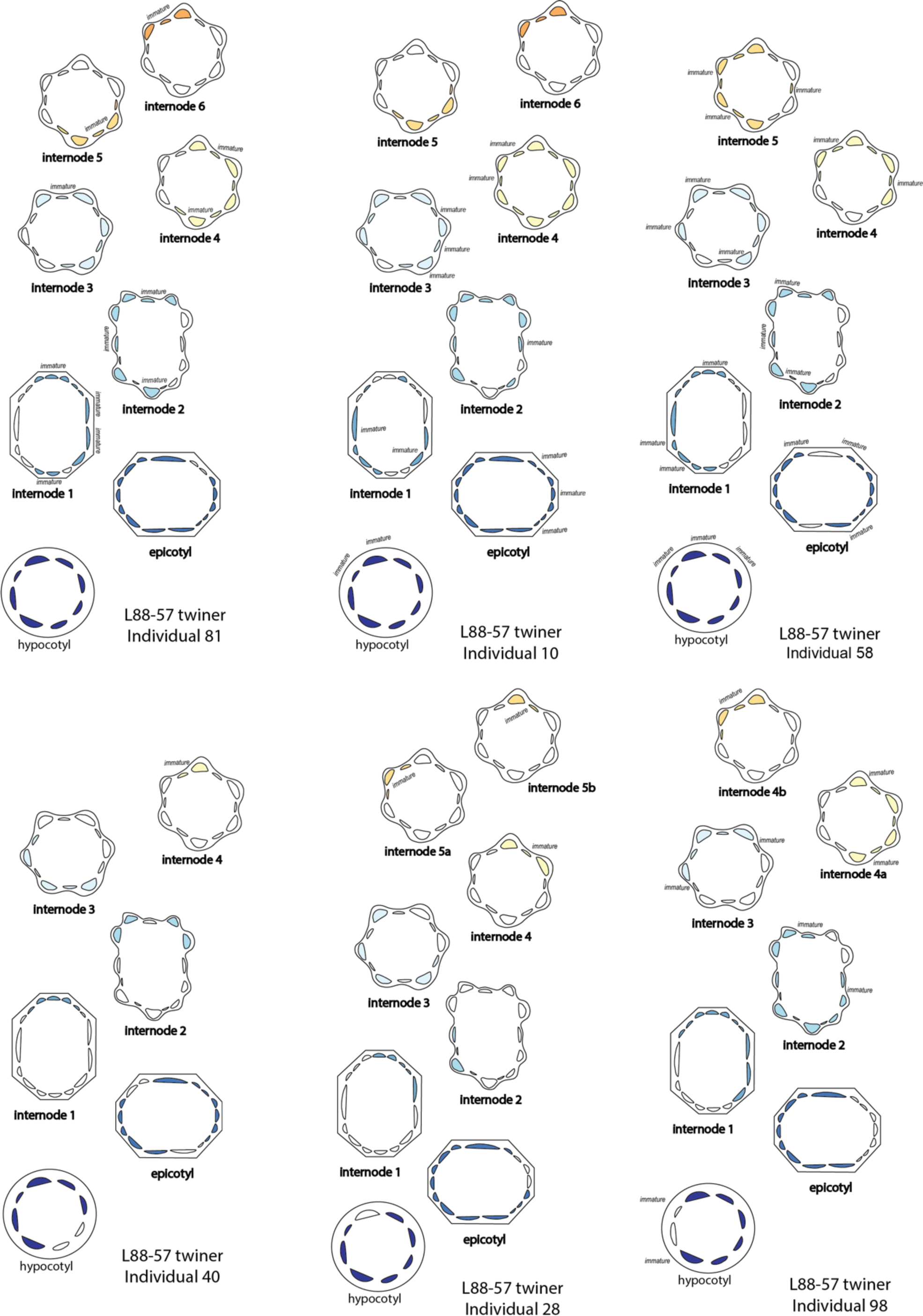
G-fiber positional data along the stems of L88-57 Twiners.

**Figure S5:**
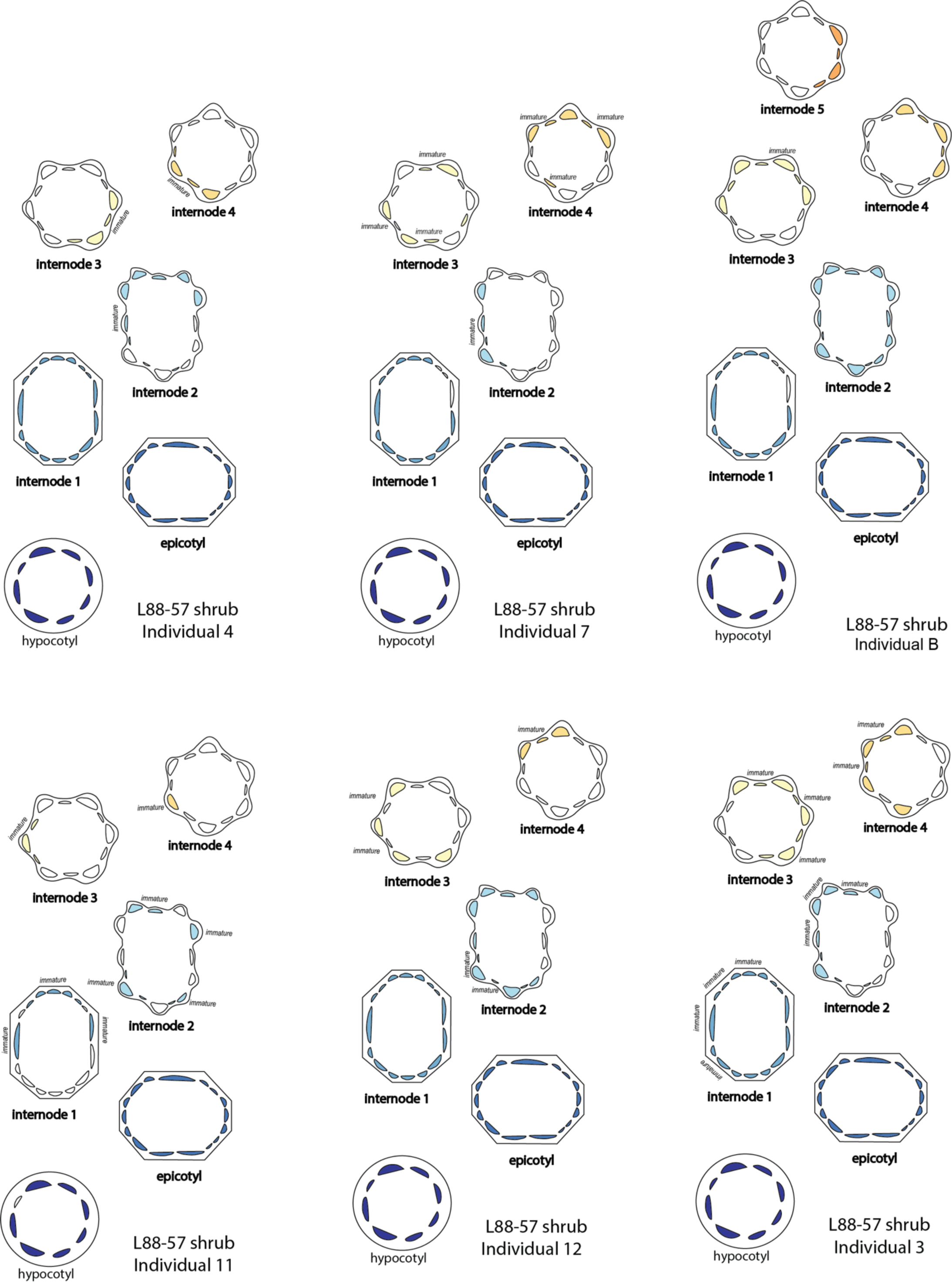
G-fiber positional data along the stems of L88-57 shrubs.

**Figure S6:**
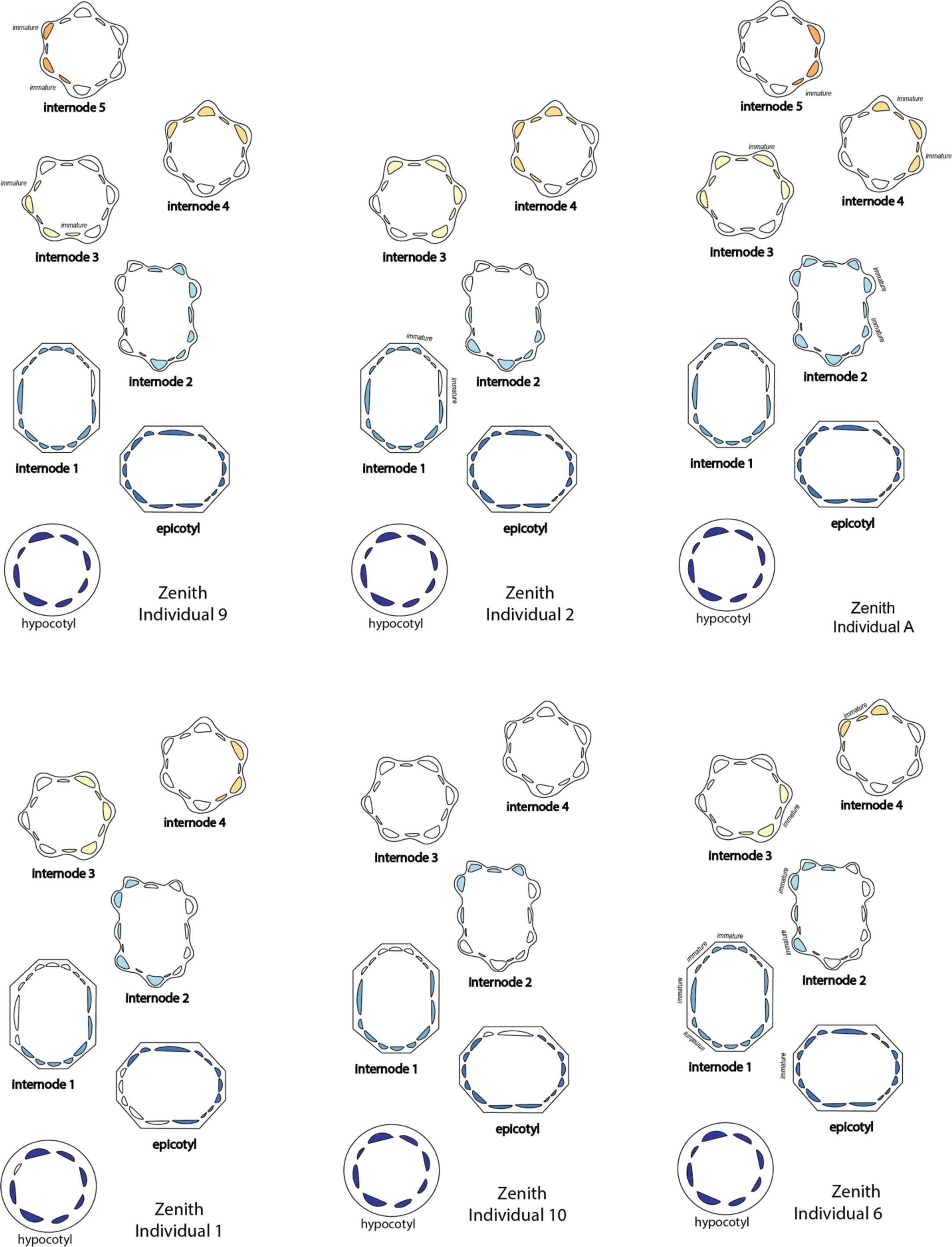
G-fiber positional data along the stems of ‘Zenith’.

**Figure S7:**
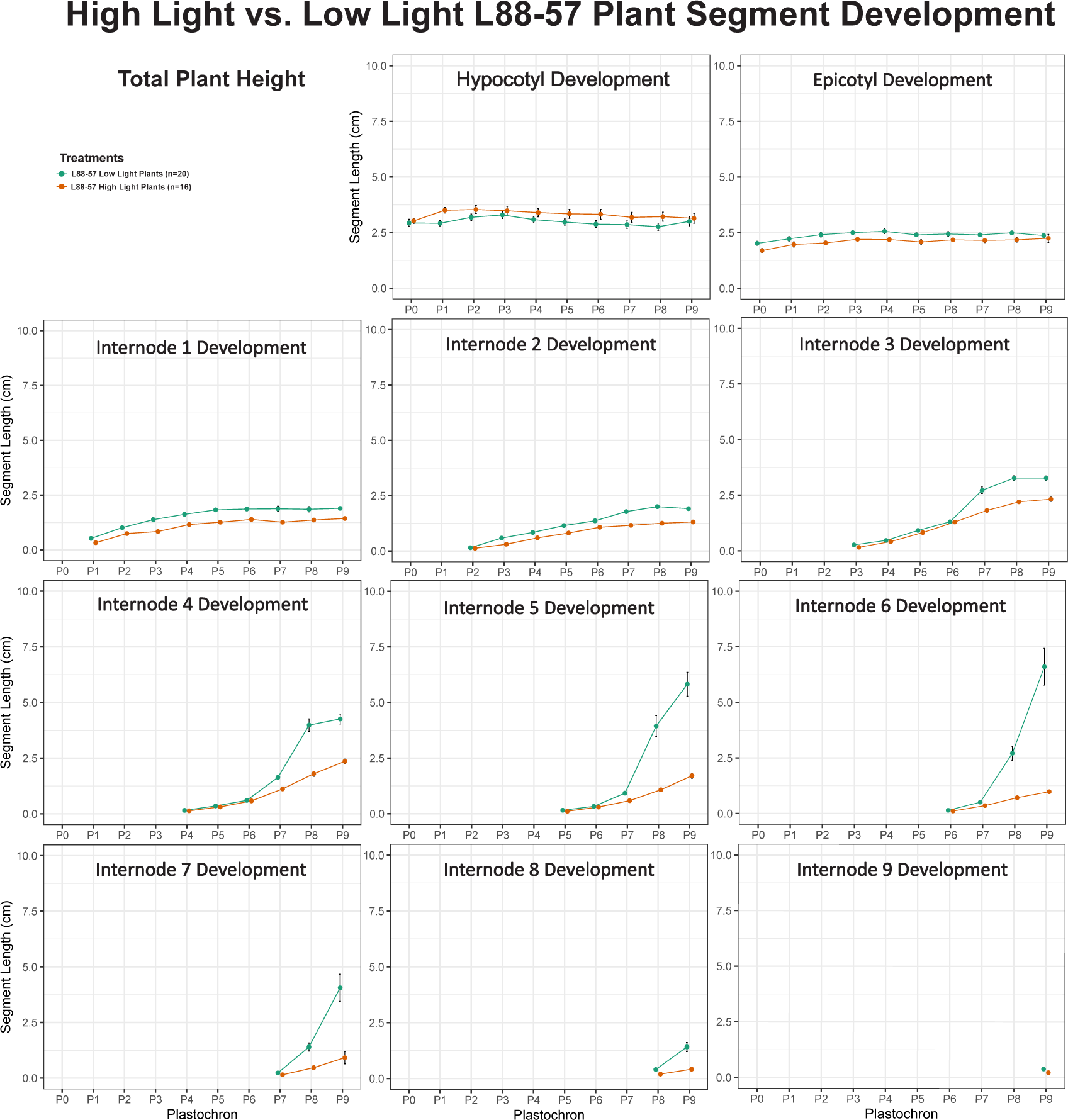
Total height and internode elongation measurements of L88-57 low light (n=20) and high light plants (n=18, except plastochron 4 and plastochron 5 has n=17) over stages of plastochron development. Plastochron 7 (P7) marks the beginning of differentiation between high vs. low light plants. Internode 3 to internode 5 of low-light plants begin elongating at P7, while high-light plants remain relatively truncated. Total height comparisons also show low-light plants differentiating from high-light plants at P7. Total height was measured by using the entire stem axis rather than adding internode measurements. Green points represent plants grown under low light (130 μmol/m2/s PAR) and orange presents plants exposed to high light (460 μmol/m2/s PAR).

**Figure S8:**
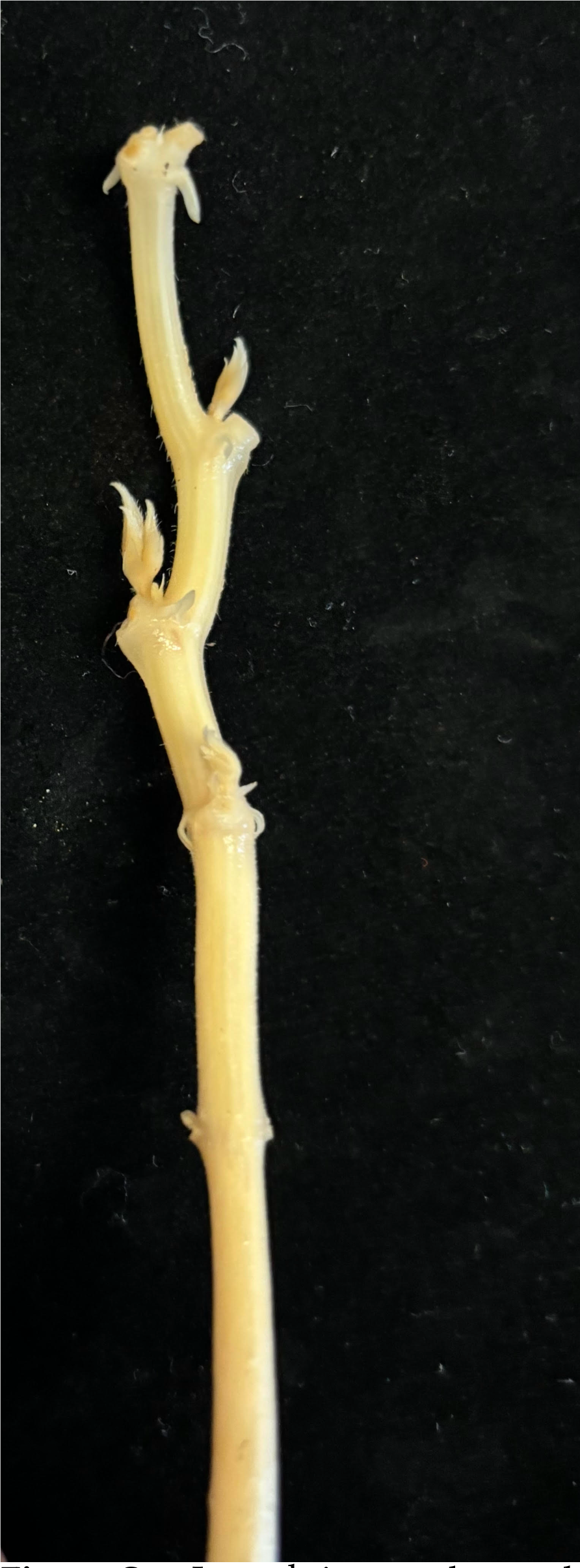
In each internode, a subtle curvature is visible, corresponding to a concave and convex side. This phenotype is present in all groups in this study (L88-57 twiners L88-57 shrubs, and Zenith shrubs). Images from L88-57 shrub, individual # 9.

